# Reaction-enabled, highly sensitive super-resolution imaging

**DOI:** 10.64898/2026.02.04.703714

**Authors:** Wenxin Zhu, Jiahui Gui, Ao Guo, Ziqing Zhang, Zhenqian Han, Weisong Zhao, Jiandong Feng

## Abstract

Super-resolution microscopy breaking the optical diffraction limit has led to success in illuminating the complexity of biology with fluorescence—light-excited luminescence^1,2^. Here, we propose another approach for super-resolution imaging with electrochemiluminescence (ECL)—reaction-excited luminescence^3,4^. While featuring high sensitivity in molecular analytics owing to its laser-free excitation, the current reaction-driven imaging suffers from limited resolution due to low photon budget^4^. Efficient ECL imaging of intracellular organelles remains elusive. To meet this challenge, we develop RIED (Reaction-based super-resolution Imaging via Entropy-weighted correlation combined with Deconvolution), which builds upon two pivotal advancements: (i) unprecedented ECL intracellular organelles imaging through a 1,000-fold enhancement in ECL emission, and (ii) the super-resolution ECL imaging by the spatiotemporal acquisition and efficient utilization of highly dynamic ECL profiles. RIED allows for the first three-dimensional super-resolution ECL imaging with a spatial resolution of ∽97 nm in the lateral dimension and ∽235 nm in the axial dimension. This allows to reveal outer mitochondrial membrane topology and distinct microtubule organizations in different mitosis phases, offering a chemistry-based imaging capability for high-throughput imaging and highly sensitive interfacial imaging, which provides a new dimension into biology.

## MAIN

Optical microscopy has been instrumental in advancing our fundamental understanding of biological systems. The use of probe molecules significantly enhances biological imaging contrast, enabling the visualization of intricate intracellular organelles and their specific functions. Consequently, fluorescence microscopy has emerged as one of the most prevalent and widely used techniques in biological imaging, which has achieved remarkable success over the past decades by surpassing the optical diffraction limit^5-8^.

Luminescence generated from chemical reactions, such as electrochemiluminescence (ECL)^3,4,9^, chemiluminescence (CL)^10,11^, and bioluminescence (BL)^12,13^, is fundamentally different from light-excited luminescence and can intrinsically eliminate laser-induced effects, featuring zero background and a high signal-to-noise ratio (SNR) that has proven practical applications in molecular diagnostics. We have recently demonstrated direct imaging of single-molecule ECL reactions^14^, an ultimate sensitivity limit for single-molecule detection. Moreover, the reaction-enabled microscopy links light emission directly to chemical reactivity, providing a unique view into the chemical heterogeneity of the biological complexity.

However, such reaction-based luminescence systems typically suffer from low photon budgets due to their inherent efficiency limitations. Consequently, state-of-the-art ECL, CL, and BL imaging have all struggled to achieve high-quality cell imaging^15-17^. Efforts to improve the performance of ECL-based cell imaging have focused on increasing the photon emission by engineering amplified probe systems or extending the imaging time^18,19^. Nevertheless, these remedies entail limitations in effective labeling and imaging resolution. As a result, in contrast to fluorescence imaging that now even aims for super-duper resolution^20,21^, the development of ECL cell imaging is a far behind process that mainly focuses on the extracellular or the whole-cell level. To date, even efficient ECL imaging of intracellular organelles has not been demonstrated, leaving such super-resolution imaging an unattainable task.

Here, we present a universal approach for super-resolution imaging with reaction-based luminescence, namely Reaction-based super-resolution Imaging via Entropy-weighted correlation combined with Deconvolution (RIED) (**Supplementary Video 1**). The development of RIED is based on two major technical breakthroughs in this work. i. We enable the first unambiguous ECL imaging of intracellular organelles by exploiting a highly efficient ECL system that improves the photon emission by 1,000-fold. ii. The enhancement of ECL emission allows to establish a unique spatiotemporal ECL recording strategy to obtain specific reaction-excited imaging information content, which is harvested in RIED to achieve unprecedented super-resolution, three-dimensions (3D), and high-throughput ECL cell imaging, demonstrating biological imaging capabilities comparable to modern super-resolution fluorescence microscopy, all while operating at zero background. We further applied RIED to visualize the microtubule networks at different mitosis phases, revealing its unparalleled sensitivity at the interface that can provide new insights complementary to super-resolution fluorescence imaging.

## RESULTS

### RIED concept

To implement the RIED concept, we first aim to address the challenge of intracellular imaging with ECL. The key experimental issues stem from inefficient labeling and low luminescence efficiency. The prevailing ECL probes struggle to specifically label intracellular organelles due to size constraints, while the limited excitation efficiency from intracellular regions further hinders sufficient luminescence. As such, conventional ECL microscopy only provides extra-cellular, whole-cell, or obscure sub-cellular information^15,16,22^. Imaging of intracellular organelles with ECL remains a challenge.

We first developed a high-efficiency ECL cell imaging system. Particularly, Bis-(2-Hydroxyethyl) aminotris (Hydroxymethyl) methane (Bis-tris) was employed as the co-reagent to replace commonly used Tripropylamine (TPrA) owing to its superior free radical diffusion distance^23^. This remarkably improves the ECL imaging performance by 1,000-fold greater than that of TPrA at the same voltage (**Extended Data Figs. 1** and **2**). More importantly, unlike using large clustered ECL emitters for luminescence amplification that can reduce labeling efficiency^18,19^, we utilized single Ru(bpy)_3_^2+^ molecules anchored to the second antibody as the luminescent probe, which exhibits excellent binding specificity and can efficiently label the target cellular structures with high coverage via immunolabelling (**Fig. 1a**). This allowed us to image microtubules, a crucial organelle for intracellular transport and maintaining cellular morphology^24^ for the first time with ECL. With the substantially enhanced ECL performance, microtubule filaments can be clearly visualized even under a 20 ms exposure time (**Supplementary Fig. 1a, 1b, Supplementary Video 2**). Through these strategies, we managed to drive ECL from “whole-cell imaging” and “extracellular imaging” to “intracellular organelle imaging” (**Supplementary Note 1**).

**Fig. 1.**
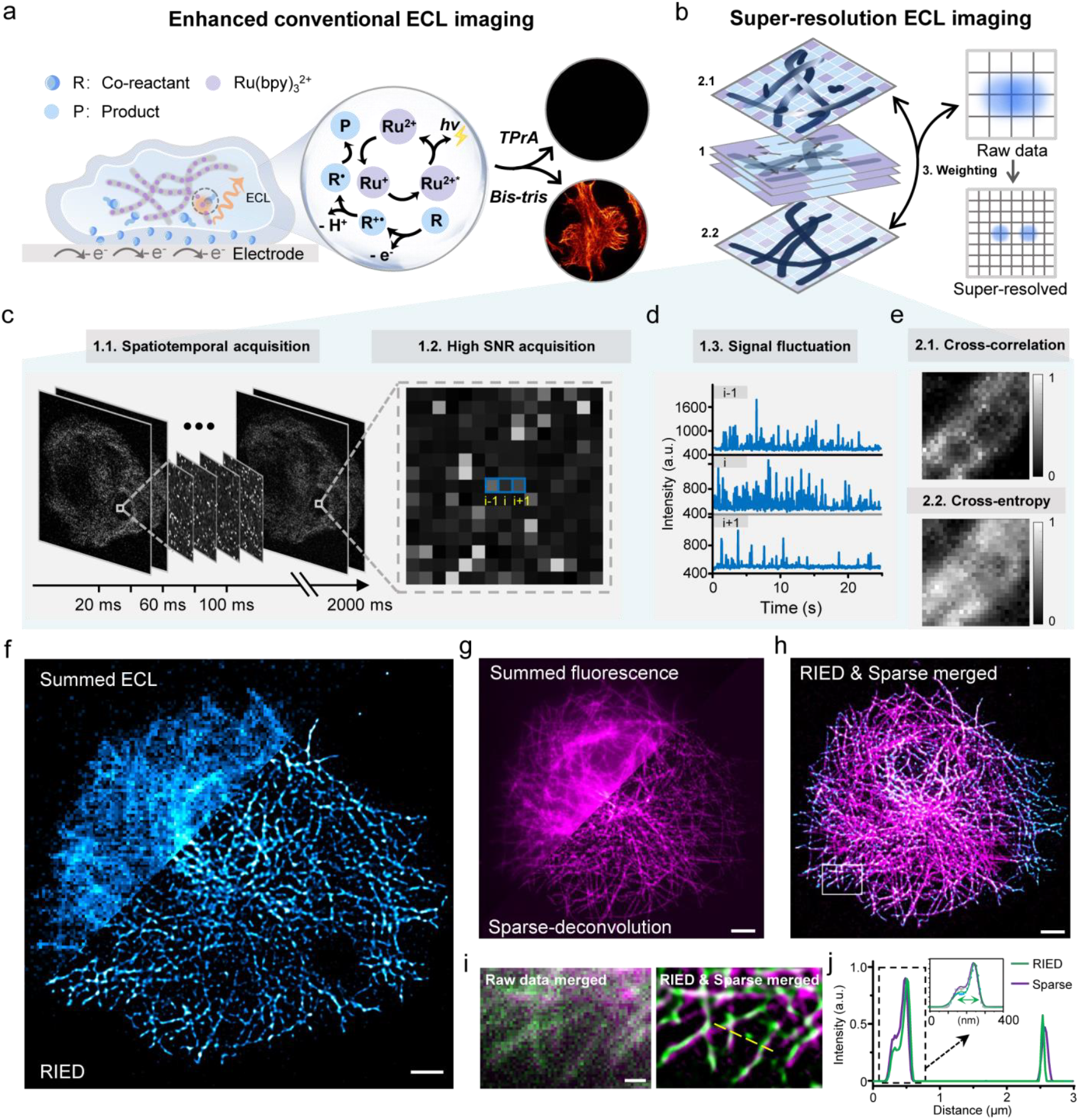
Super-resolution ECL imaging with RIED. **a**, Schematic of ECL excitation and emission for imaging Ru(bpy)_3_^2+^ labeled intracellular microtubule filaments with different co-reactant (TPrA and Bis-tris). **b**, Schematic of core algorithms used in RIED. **c**, High-speed acquisition (100 frames, 20 ms per frame) of ECL stacks with spatiotemporal separation properties. **d**, Intensity profiles of three separate individual pixels highlighted by blue lines in ‘c’. **e**, Spatiotemporal correlation mapping of zoomed view from ‘c’ for super-resolution imaging (top). The imaging SNR and structural integrity are further improved by the cross-entropy coefficient map (bottom). **f**, ECL image of labeled microtubules before (top left) and after (bottom right) RIED reconstruction. Scale bar, 5 µm. **g**, Corresponding fluorescence images of the microtubules in ‘f’ before (top left) and after Sparse deconvolution (bottom right). Scale bar, 5 µm. **h**, The merged image of RIED and fluorescence Sparse deconvolution. Scale bar, 5 µm. **i**, Zoomed views from the white box in ‘h’ before (left) and after (right) super-resolution reconstruction. Scale bar, 500 nm. **j**, Intensity profiles and multiple Gaussian fitting of RIED and Sparse deconvolution for microtubule filaments indicated by yellow dashed line in ‘i’.

The second task is how to overcome the super-resolution challenge for such reaction-based luminescence. Although the ECL probe is labeled to cellular structures suppressing diffusion, the spatial resolution is still limited by the diffraction. Mainstream ECL microscopy commonly extends the exposure time to tens of seconds to accumulate a single image, overlooking the rich spatiotemporal information. However, the reaction-based luminescence is a highly dynamic process, intrinsically triggered by individual reaction events. By recording these spatiotemporally isolated ECL processes, one may be able to extract the distinct reaction-based profiles and utilize them to achieve super-resolution imaging (**Fig. 1b, Extended Data Fig. 1**). Thanks to the enhanced ECL emission, we can perform the spatiotemporal acquisition of transient luminescence events in an image sequence, which reveals the detailed features of ECL imaging such as zero-background, high SNR, and intensity fluctuations (**Fig. 1c, 1d, Extended Data Fig. 3**). We also noticed significant changes in ECL intensity and emission rate during the reaction (**Extended Data Fig. 3c**), reflecting a unique reaction-exited profile. We then designed an algorithm for super-resolution ECL imaging by leveraging these features.

In principle, the ECL emission exhibits high-contrast blinking of independent emitters, and we can exploit these fluctuations to effectively enhance the spatial resolution. Based on the ECL events recorded over time, we performed spatiotemporal cross-correlation analysis to sequences of photon events for excluding pixels dominated by overlapping emitters and narrowing the point spread function (PSF) width (**Fig. 1b, 1e, Supplementary Fig. 1c, Methods**), drawing on our recently developed fluorescence super-resolution method^25^. However, the ECL imaging exhibits inherent spatial heterogeneity from the emitter molecules, leading to a structural discontinuity. To address this, we calculated an entropy map on a sub-pixel basis, before cross-correlation analysis. This map quantifies the expected photon information content and serves as a weighting factor to compensate for the correlation cumulant affected by the spatiotemporal ECL heterogeneity (**Fig. 1b, 1e**). Besides, the weighting of the entropy map can also highlight the ECL contrast, despite low photon budget conditions (**Supplementary Fig. 1c**). Finally, a deconvolution step was introduced for spatial resolution maximization and image quality refinement (detailed workflow in **Extended Data Fig. 4, Methods**).

Taken together, we have here established an ECL-based super-resolution cell imaging method—RIED. Tailored specifically for reaction-based luminescence, it demonstrates superior reconstruction performance compared with conventional super-resolution algorithms that are designed for processing fluorescence data (**Extended Data Fig. 5**)^25-28^. The development of RIED revolutionizes the visualization of intracellular organelle structures that are invisible by conventional ECL microscopy (**Fig. 1a, Extended Data Figs. 1, 2a**). For cross-validation, we compared the reconstructed RIED results with super-resolution fluorescence imaging data^29^, both targeting microtubules of the same cell, and their composite image exhibits a high consistency (**Fig. 1f-1h, Supplementary Fig. 2a, 2b**). By analyzing the overlapped regions, both visual inspection and intensity profiles of the reconstructed microtubules correspond well (**Fig. 1i, 1j**), further confirming the reliability of RIED for intracellular imaging at the sub-diffraction scale. Note in ECL imaging, in contrast to fluorescence imaging, the unique excitation mechanism imparts its surface sensitivity and high SNR as the reaction is triggered at the electrode surface^16^ and no excitation light is introduced **(Supplementary Fig. 3, Supplementary Note 2)**. Therefore, our RIED allows for the distinct observation of microtubule filaments near the electrode surface (**Fig. 1h**).

### Resolvability of RIED

We next evaluated the spatial resolution of RIED (**Fig. 2a-2c, Supplementary Fig. 2c**). Compared to conventional ECL imaging typically at sub-micrometer resolution, RIED shows a remarkable improvement in spatial resolution, reaching 97 nm as determined by Fourier ring correlation (FRC) analysis (**Fig. 2d**)^30^. The peak-to-peak separation of the intertwining microtubule filaments in **Fig. 2c** suggests that the resolving power of RIED (**Fig. 2e**) is comparable to super-resolution fluorescence microscopy^25^. Beyond the cytoskeleton, we also looked at the mitochondria with RIED, which can be clearly reconstructed with a FRC resolution of 121 nm (**Fig. 2f, Supplementary Fig. 2d**). Interestingly, various forms of outer mitochondrial membrane, e.g., tubular, spheroid, small vesicles, and fragments, can be further distinguished (**Fig. 2g**). This imaging capability marks a super-resolution leap over conventional ECL cell imaging, which lacks the resolvability for dissecting sub-cellular structures.

**Fig. 2.**
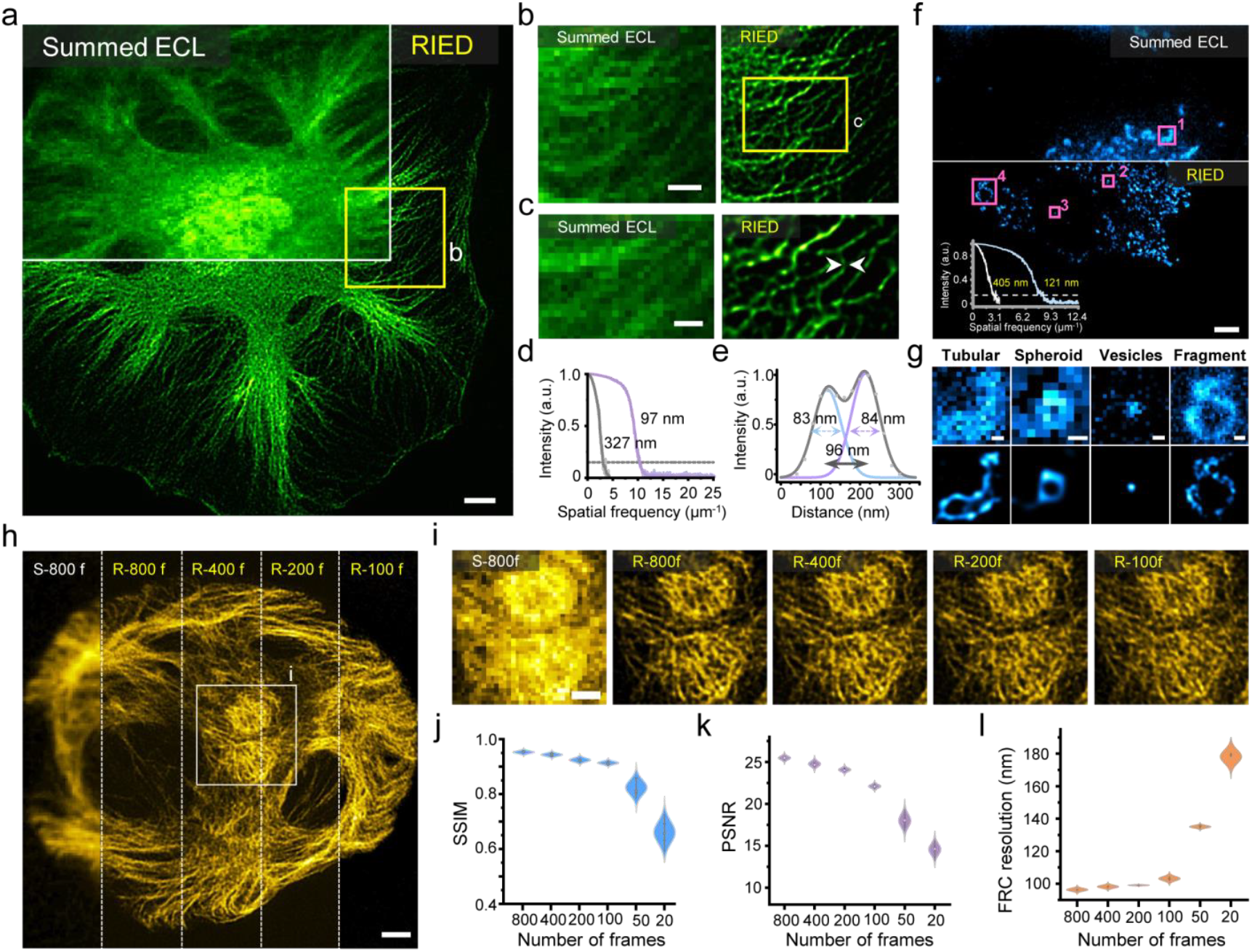
Spatiotemporal resolution of RIED. **a**, Microtubule filaments in a HeLa cell imaged by summed ECL (top left) and RIED (bottom right). Scale bar, 5 µm. **b**, Magnified views from the yellow box in ‘a’. Scale bar, 3 µm. **c**, Zoomed views from the yellow box in ‘b’. Scale bar, 1 µm. **d**, FRC analysis of the summed ECL and RIED images in ‘a’. **e**, Intensity profiles and multiple Gaussian fitting of the reconstructed microtubule filaments indicated by the white arrows in ‘c’. **f**, Mitochondria in a COS-7 cell imaged by summed ECL microscopy (top) and RIED (bottom). Scale bar, 5 µm. Insert: FRC analysis of the summed ECL and RIED images. **g**, Enlarged views of summed ECL (top) and RIED images (bottom) from the red boxes in ‘f’. Scale bars, 500 nm. **h**, From left to right: Microtubule filaments in a HeLa cell under 800-frame summed ECL (S-800 f) and RIED images reconstructed with 800 (R-800 f), 400 (R-400 f), 200 (R-200 f), and 100-frames (R-100 f). Scale bar, 5 µm. **i**, Zoomed views from the white box in ‘h’. Scale bar, 1 µm. **j**,**k**,**l**, SSIM (**j**), PSNR (**k**) and FRC (**l**) analysis of images in ‘h’. Each distribution is obtained from 10 parallel measurements.

While improving spatial resolution typically requires prolonged exposure, RIED can efficiently harness each ECL photon to generate a super-resolution image from just 100 raw frames at 20 ms exposure, achieving unprecedented temporal resolution in ECL imaging. By benchmarking the RIED results from different numbers of frames (**Fig. 2h, 2i**), we found that 100-frame is sufficient for high-quality super-resolution reconstruction, which is confirmed by visual inspection, structure similarity index measure (SSIM>0.9), peak signal-to-noise ratio (PSNR>20), and FRC analysis (<110 nm) (**Fig. 2j-2l, Supplementary Fig. 4** and **5**). We also found that the temporally interpolated data yield better reconstruction quality than raw data (**Extended Data Fig. 6, Methods**).

### 3D super-resolution ECL imaging

As organelles display complex 3D architectures in cells, dissecting their 3D distributions will certainly contribute to a deeper understanding of their functions and variations. However, up to now, 3D ECL microscopy remains a technical challenge, primarily due to difficulties in regulating the ECL excitation depth. In our experiments, we observed a significant change in the ECL emission depth which directly responds to the applied voltage amplitude (**Fig. 3a**). At 1.0 V, the emission predominantly occurs near the surface of the electrode, and when increasing the voltage to 1.8 V, the center of the cell gradually exhibits more pronounced photon emission (**Fig. 3b**). Leveraging this phenomenon, we further investigated the capacity of ECL for 3D image reconstruction. By synchronizing voltage control with changes in the imaging focal plane, the 3D-RIED can clearly distinguish the cytoskeleton network in all dimensions (**Fig. 3c-e, Supplementary Fig. 6, Supplementary Video 3**). Notably, microtubules spaced ∽240 nm apart in the axial axes are clearly resolved with 3D-RIED (**Fig. 3d**), and the lateral resolution is ∽116 nm (**Fig. 3f**). The resolution in all three axes is comparable to the 3D super-resolution fluorescence imaging^25,31^, suggesting that ECL imaging can successfully capture the orientation of microtubules (**Fig. 3g**). In addition, the lateral FRC resolution in different axial cross-sections is uniformly distributed between 97 nm and 118 nm (**Fig. 3h**). Together, we demonstrated the first 3D super-resolution capability of ECL imaging, extending its application to record complex organelle networks.

**Fig. 3.**
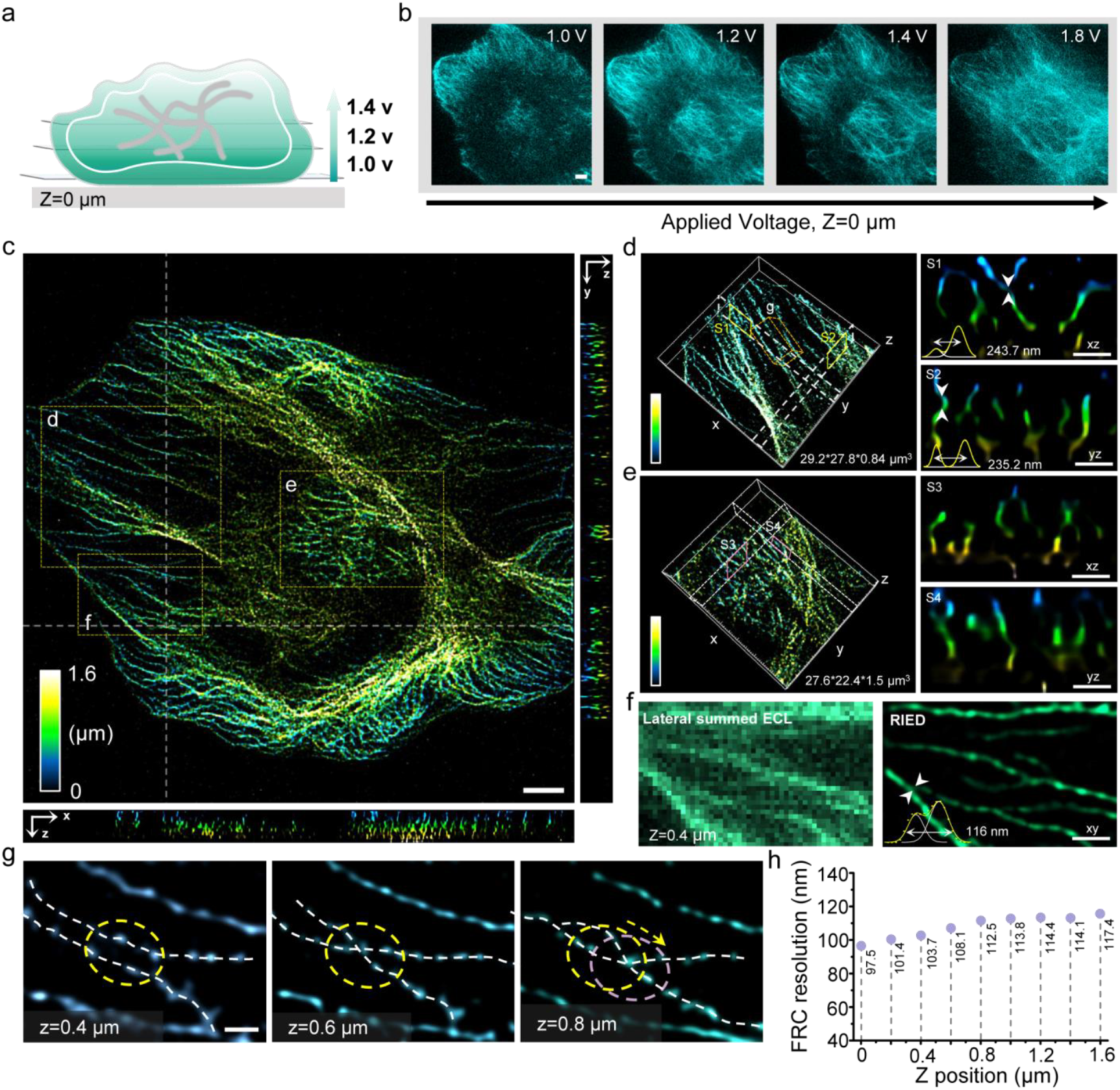
3D super-resolution ECL imaging. **a**, Schematic of the ECL excitation depth related to the applied voltage. **b**, Representative ECL images of labeled microtubules at different applied voltages. Scale bar: 5 µm. **c**, Color-coded 3D distributions of microtubules in a HeLa cell under RIED as well as the x-z and y-z cross-sections along the gray dash line, Scale bar: 5 µm. **d**, Magnified 3D rendering of the boxed region labeled ‘d’ in ‘c’ recorded by RIED. S1, S2, the x-z, y-z cross-sections along the corresponding yellow dashed boxes. The white arrows indicate the positions where the intensity profiles are taken, and the distances between peaks are obtained with multiple Gaussian fitting. Scale bars: 500 nm. **e**, Magnified 3D rendering of the boxed region labeled ‘e’ in ‘c’ taken by RIED. S3, S4, the corresponding x-z and y-z cross-sections along the pink dashed boxes. Scale bars: 500 nm. **f**, 2D zoomed lateral views from the yellow dashed box labeled ‘f’ in ‘c’ imaged by summed ECL microscopy and RIED. Scale bar: 1 µm. **g**, Lateral variations of tubulin distribution at different z-axis height from the cubic region labeled ‘g’ in ‘d’. Scale bar: 1 µm. **h**, FRC resolutions at different axial positions.

### RIED permits high-throughput imaging and analysis

We then explored RIED’s unique capabilities for cell imaging, starting with high-throughput imaging enabled by its exceptional temporal resolution. As a proof-of-concept demonstration, we imaged the intracellular microtubules on a large scale (0.53 mm × 0.53 mm) area with 3 × 3 field-of-view (FOVs) (512 × 512 pixels) each (**Fig. 4a, Supplementary Fig. 7a**) in which the microtubule filaments are clearly distinguishable (**Fig.4b, Supplementary Fig. 7b, Supplementary Video 4**). The mean rFRC resolution^30^ of full FOV is ∽144 nm (**Fig. 4c, 4d**), indicating the imaging resolution is not affected by scaling up the FOV. These analyses highlight RIED as a versatile tool for high-throughput super-resolution imaging.

**Fig. 4.**
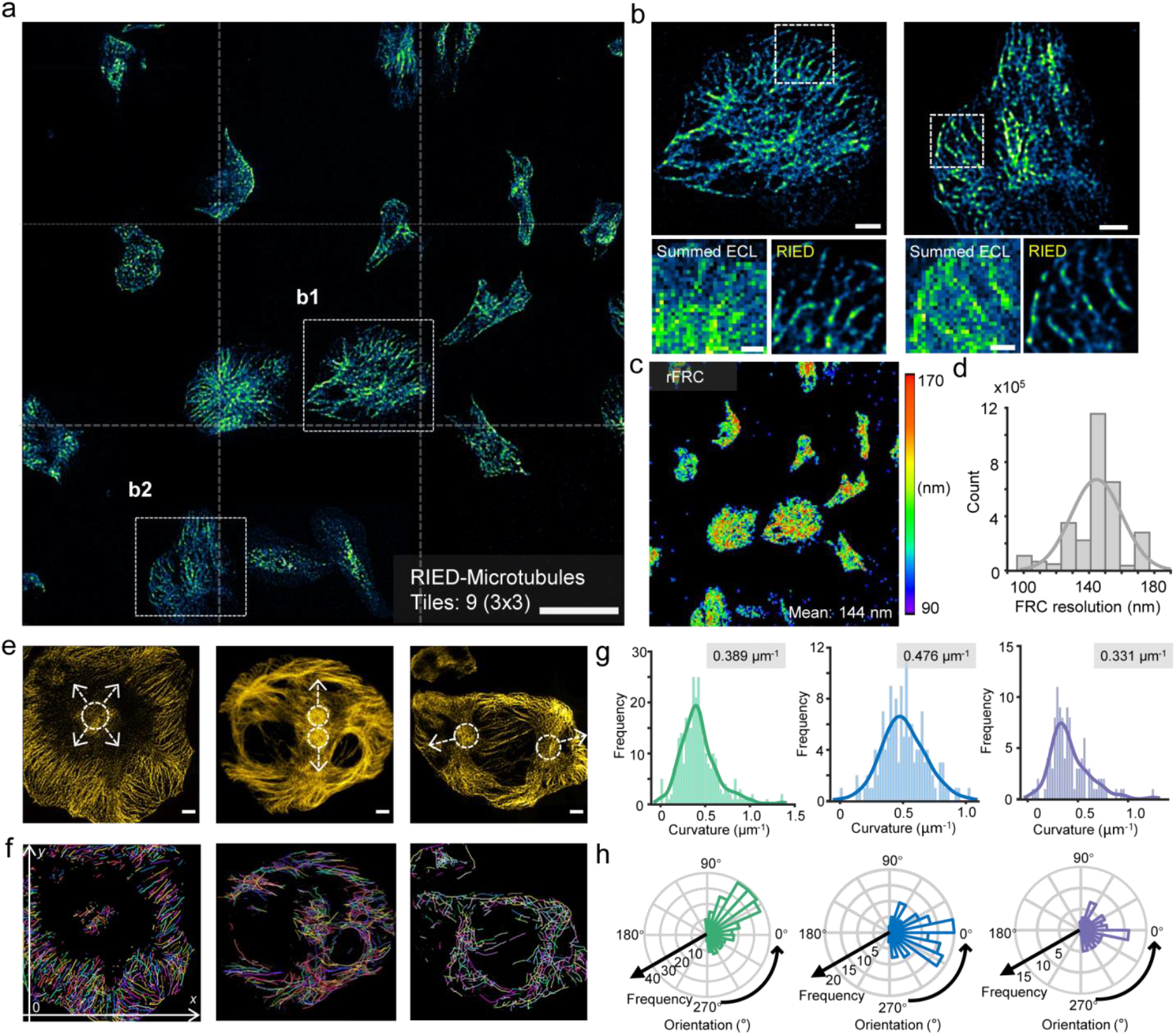
High-throughput imaging and analysis of microtubules. **a**, RIED for high-throughput super-resolution imaging of a 0.53 mm × 0.53 mm area. Scale bar, 40 µm. **b**, Enlarged microtubule distributions in a HeLa cell labeled as ‘b1’, ‘b2’ in ‘a’ (top). Scale bars, 5 µm. Summed ECL and RIED results for the area highlighted by the white dashed box in ‘b’ (bottom). Scale bars, 500 nm. **c**, rFRC mapping of ‘a’. **d**, Distribution of rFRC resolution in ‘c’. **e**, Microtubule filaments at different mitosis phases imaged by RIED. Scale bars: 5 µm. **f**, Segmented microtubule skeletons from ‘e’. **g**, Curvature distributions of microtubule filaments in ‘f’. **h**, Orientation analysis of microtubule filaments in ‘f’.

Cell division is essential for the growth, development, and reproduction of organisms, in which microtubule filaments play a crucial role by positioning and separating chromosomes^32^. Thus, we further investigated the morphology of microtubule filaments, taking advantage of the high interfacial sensitivity of RIED. According to the RIED results, the distribution of microtubules varies significantly across different phases of mitosis (**Fig. 4e**). By extracting the cytoskeleton distributions to obtain their spatial coordinates (**Fig. 4f**), we statistically analyzed microtubule filaments’ curvature and orientation across different mitosis phases. Both ECL (**Fig. 4g, 4h**) and fluorescence (**Extended Data Fig. 7**) results reveal similar mitotic dynamics: microtubule filament curvatures exhibit a biphasic pattern (initial increase followed by decrease), while their orientation gradually converges toward 0° during mitosis. This result aligns with the phenomenon that the microtubules first converge towards the poles and then adopt a straight, pole-oriented arrangement during the spindle self-assembly process^33^. Interestingly, in the same mitosis stage, the ECL result exhibits smaller curvature values and more concentrated orientation distributions than the fluorescence result. This is attributed to the interfacial sensitivity of the ECL, which takes more linear microtubules near the surface (**Fig. 4e**) into account.

### RIED unlocks highly sensitive imaging

Stemming from the unique ECL mechanism, light emission is triggered only at the electrode interface and requires no external light source. Thus, interference from scattered light or autofluorescence is minimized. These advantages of ECL significantly contribute to single-cell immune profiling^34^. The interfacial sensitivity is advantageous for imaging membrane-proximal targets such as integrin, collecting unique information complementary to fluorescence imaging (**Extended Data Fig. 8a**). The high imaging SNR is also beneficial for imaging proteins with low expression such as carcinoembryonic antigen (CEA) — a biomarker, demonstrating an enhanced detection sensitivity both in positive and negative cells than that of fluorescence super-resolution imaging (**Extended Data Fig. 8b-8e, Supplementary Note 2**). These findings highlight the high sensitivity of ECL imaging, which can provide a complementary view to fluorescence imaging.

## DISCUSSION

Our approach outperforms conventional ECL microscopy in providing both unprecedented cellular imaging capability and super-resolution capability. We established a spatiotemporally isolated ECL recording strategy based on an experimental design that enhances ECL emission by 1,000-fold. This enables systematic analysis of isolated ECL imaging stacks, revealing two key features of this reaction-excited luminescence: (i) dynamic emission fluctuations against a zero-background regime, and (ii) a time-related reactivity heterogeneity. Harvesting these highly dynamic luminescence characteristics, we designed a method to achieve super-resolution, delivering a high spatial resolution comparable to modern super-resolution fluorescence microscopy. Notably, voltage-modulated reaction excitation introduces a depth-regulated emission control— a phenomenon we exploited to demonstrate the first 3D super-resolution ECL imaging. Moreover, RIED’s high-throughput capability facilitates rapid data acquisition of large-scale samples, accelerating research on cellular heterogeneity^35^. This can extend the applications of ECL to larger-volume biological subjects, including tissue-scale super-resolution imaging.

While RIED’s performance now approaches fluorescence-based super-resolution imaging, a fundamental difference lies in their distinct excitation mechanisms. As a laser-free method featuring zero background, ECL has been successfully applied in molecular diagnostics owing to its superior sensitivity. Now adding new spatiotemporal dimensions, RIED may advance the bioassay field to the desired single-cell immune profiling. In addition, the voltage-driven excitation profiles of ECL pronounced its surface sensitivity, enabling ultrasensitive detection of cellular proximal analytes.

Given the basic principle of RIED, it is universally applicable to other reaction-based luminescence imaging systems, including CL microscopy and BL microscopy. The potential integration of CL and BL can further extend the super-resolution imaging capabilities to drive diverse applications, such as long-term observation of living cells and *in vivo* bioimaging due to its good biocompatibility^36^. Unlike conventional optical microscopy relying on optical system optimization, this methodology manipulates the imaging capability by tailoring the luminescence reaction profile, meaning that it can flexibly design reaction systems according to different application scenarios^37^.

Finally, the image contrast in reaction-enabled microscopy is driven by chemical reactivity, which intrinsically visualizes the activity of the molecule rather than just molecular positions. This unique mechanism may revolutionize biological imaging by providing an additional activity view dimension, empowering direct functional analysis such as cell redox imaging. Thus, we anticipate that the development of RIED will highlight a new imaging paradigm, i.e., chemical reaction-based super-resolution microscopy, which may unlock unprecedented capabilities to uncover chemistry-related information in biological and chemical systems.

## Data availability

All data that support the findings of this study are available from the corresponding author upon request.

## Code availability

The code to perform RIED super-resolution reconstruction will also be available at https://github.com/SR-Wiki/RIEDm, upon final publication.

## Acknowledgments

This work was funded by the National Natural Science Foundation of China (grant numbers: 21974123 to J.F., 32422052 and 62305083 to W-S.Z.), the National Key R&D Program of China (grant numbers: 2020YFA0211200 to J.F., 2022YFC3400600 to W-S.Z.), the Fundamental Research Funds for the Central Universities (grant number: K20220088), and the Young Elite Scientists Sponsorship Program by China Association for Science and Technology (grant number: 2023QNRC001 to W-S.Z.). J.F. acknowledges the support from the New Cornerstone Science Foundation through the XPLORER Prize. W-S.Z. acknowledges the facility support from the Frontiers Science Center for Matter Behave in Space Environment, Harbin Institute of Technology. We thank the Micro and Nano Fabrication Platform at Zhejiang University for facility support.

## Author Contributions

J.F. and W-X.Z. initiated the project and developed the imaging methodology; G.J. and W-S.Z. developed the super-resolution algorithm; W-X.Z., A.G. and Z.Z. performed the experiments; W-X.Z., G.J., and Z.H. analysed the data; J.F. and W-S.Z. conceived and supervised the study; W-X.Z., G.J., W-S.Z., and J.F. wrote the manuscript with input from all authors. All authors participated in the discussions and data interpretation.

---

**Extended Data Fig. 1.**
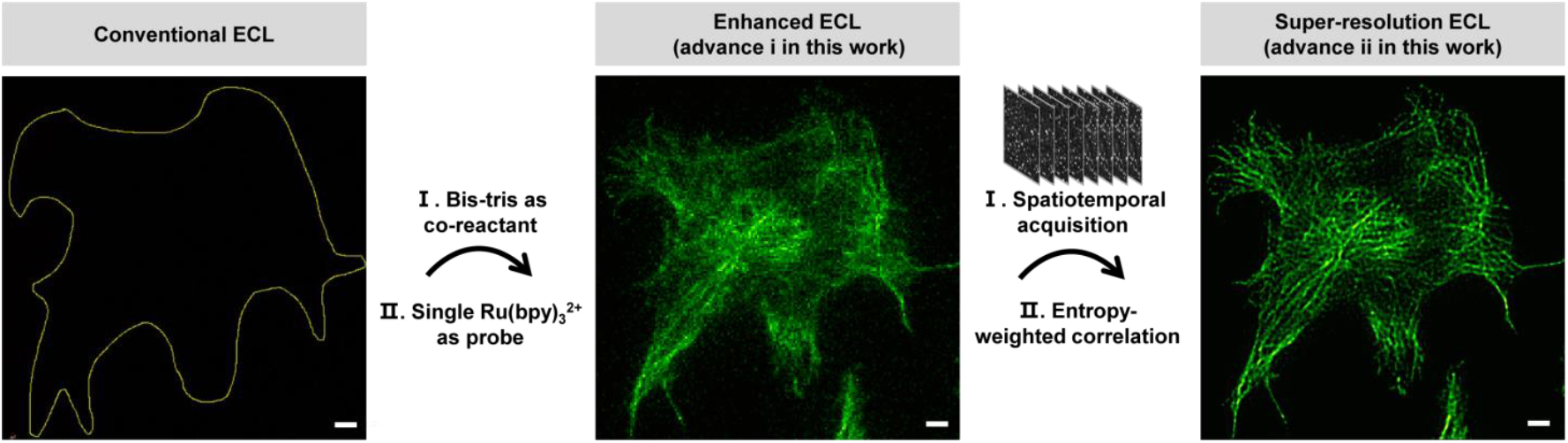
Two-stage developments for fulfilling super-resolution ECL cell imaging. The first microtubule imaging with ECL is achieved here by introducing Bis-tris as the co-reactant and a single Ru(bpy)_3_^2+^ molecule as the labeling probe. Super-resolution imaging of microtubule filaments with ECL is realized by analyzing the entropy-weighted cross-correlation of spatiotemporally acquired ECL events. Scale bars: 5 µm.

**Extended Data Fig. 2.**
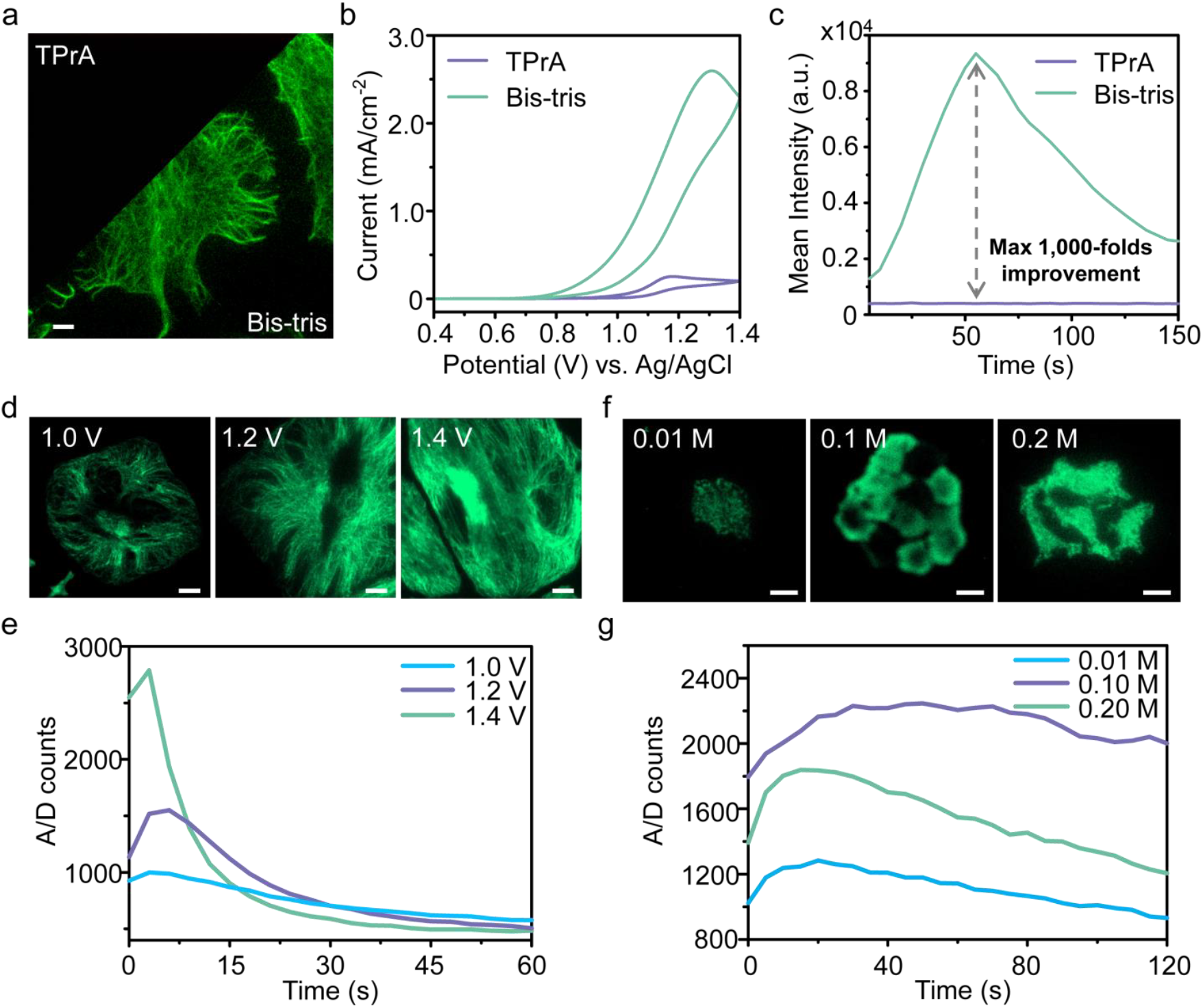
Optimization of ECL imaging conditions for intracellular observation. **a**, Summed ECL image of Ru(bpy)_3_^2+^ labeled microtubule filaments with TPrA (top left) and Bis-tris (bottom right) as co-reactant respectively. Concentration of co-reactant: 100 mM. Scale bar: 5 µm. EM Gain: 500, exposure time: 5 s. **b**, CVs of TPrA and Bis-tris as co-reactant. Solution: 100 mM TPrA or 100 mM Bis-tris with 0.01 M PBS. ITO diameter: 10 mm. Scan rate: 100 mV/s. **c**, Intensity profile of summed ECL images with Bis-tris and TPrA as co-reactant. EM Gain: 500, exposure time: 5 s. The intensity improvement is calculated as the ratio of the respective signal intensity minus the background intensity (450 analogue-to-digital counts). **d**, Summed ECL images under the potential of 1.0 V, 1. 2 V, and 1.4 V (*vs*. Ag/AgCl). Scale bars: 5 µm. EM Gain: 500, exposure time: 3 s. **e**, Time trace of the corresponding detected analogue-to-digital (A/D) counts of the cells in ‘d’. **f**, Summed ECL images with different concentrations of bis-tris under 1. 2 V (*vs*. Ag/AgCl). Scale bars: 30 µm. EM Gain: 500, exposure time: 1 s. **g**, Time trace of the corresponding detected A/D counts of the cells in ‘f’.

**Extended Data Fig. 3.**
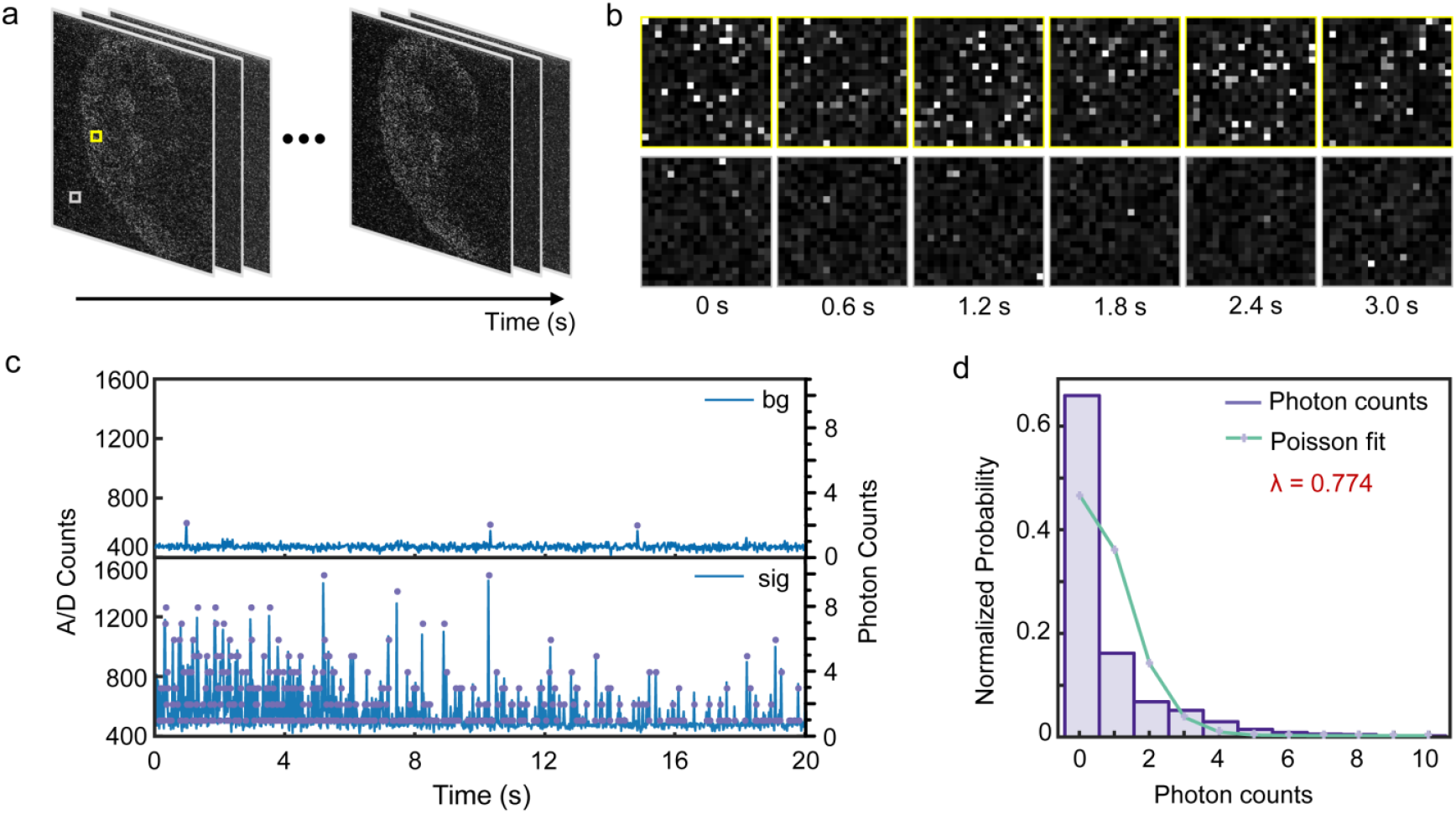
Spatiotemporal acquisition of ECL signals. **a**, Representative frame trace under an exposure time of 20 ms, EM gain: 500. **b**, Magnified snapshots of ECL signal and background labeled with yellow and gray boxes in ‘a’. **c**, Time traces of the detected analogue-to-digital (A/D) counts and photon counts of single pixel in ‘a’. **d**, Photon count distributions, and the Poisson fit in ‘c’.

**Extended Data Fig. 4.**
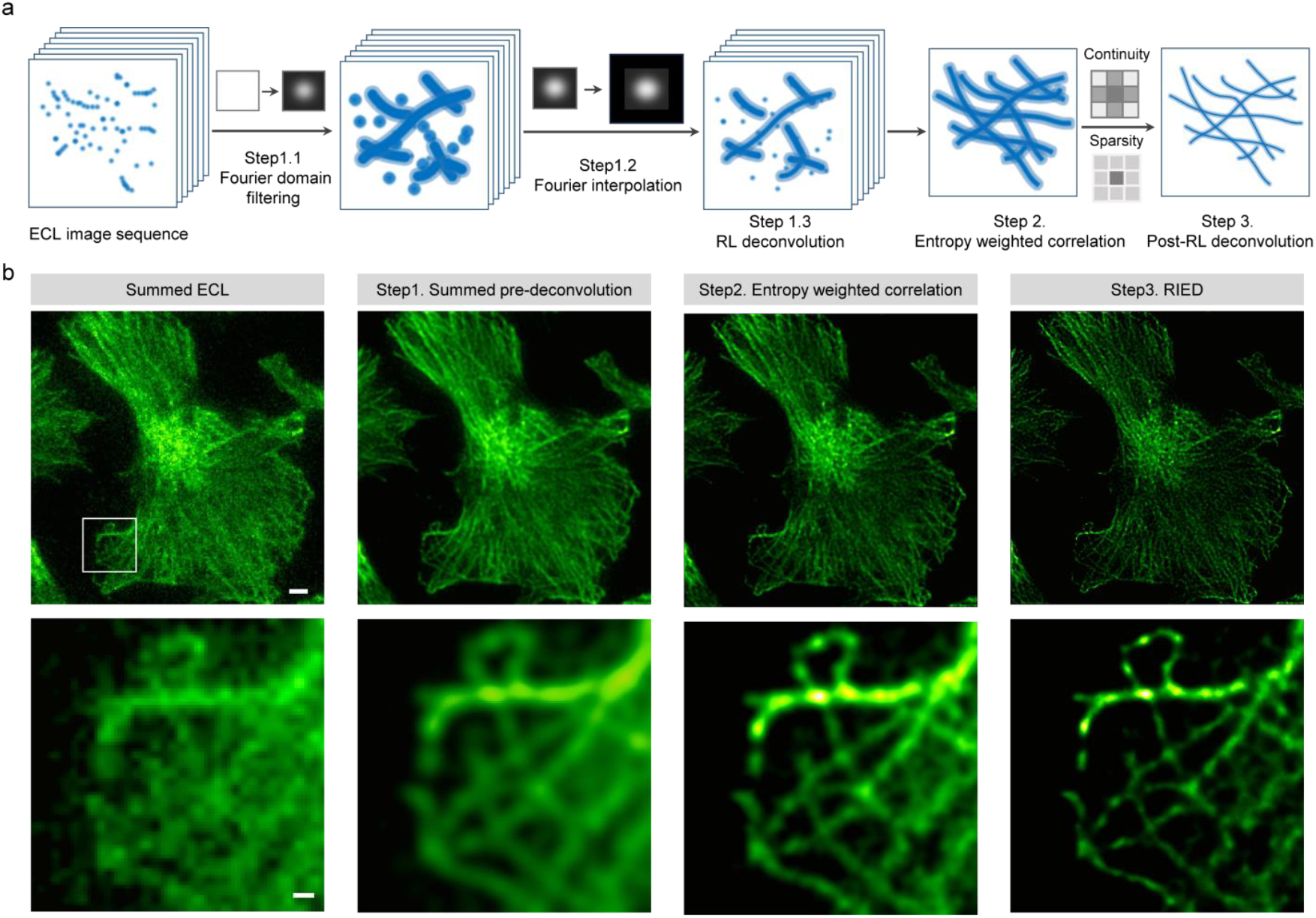
Computational workflow of RIED. **a**, Illustration of the workflow in RIED reconstruction. **b**, Detail reconstructed images after each process step in ‘a’. Scale bars: 5 µm (top image), 1 µm (bottom image).

**Extended Data Fig. 5.**
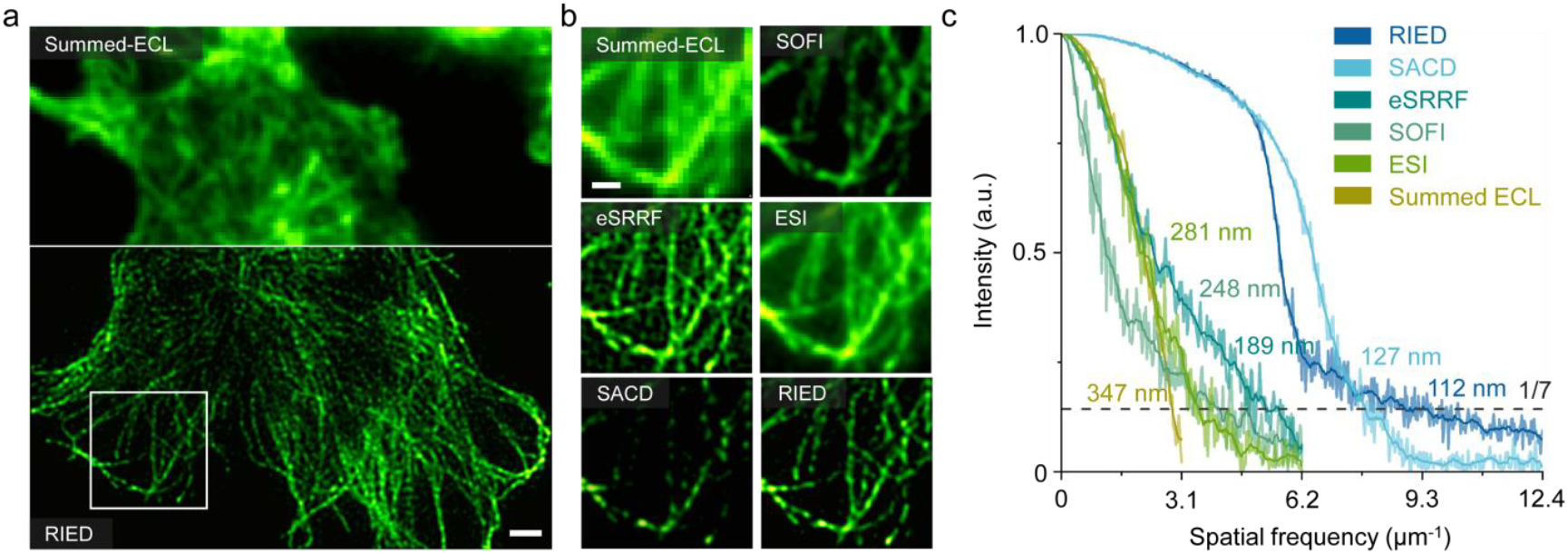
Reconstruction comparisons of RIED with other fluorescence super-resolution methods. **a**, Summed ECL image (top) and RIED image (bottom). Scale bar: 5 µm. **b**, Enlarged views of the white box in ‘a’ processed by different methods. Scale bar: 3 µm. **c**, FRC analysis of reconstructed images with different methods.

**Extended Data Fig. 6.**
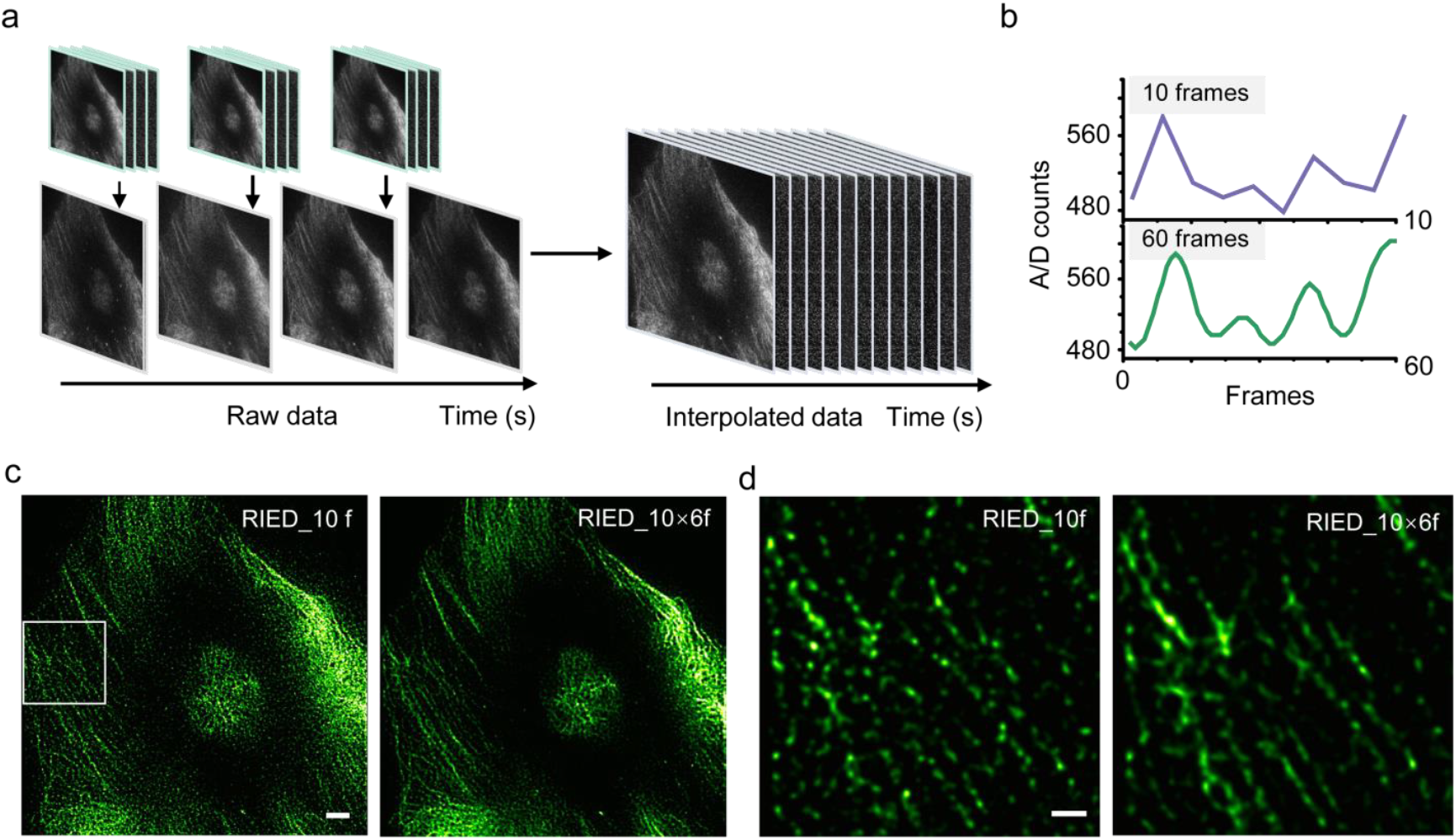
Timeline interpolation in RIED. **a**, Illustration of the interpolated raw data for RIED reconstruction. **b**, Intensity profiles of the raw data and interpolated data. **c**, RIED reconstruction result from raw data (10 frames) and time-interpolated data (virtual 60 frames). Scale bar: 5 µm. **d**, Enlarged microtubules of the white box in ‘c’. Scale bar: 2 µm.

**Extended Data Fig. 7.**
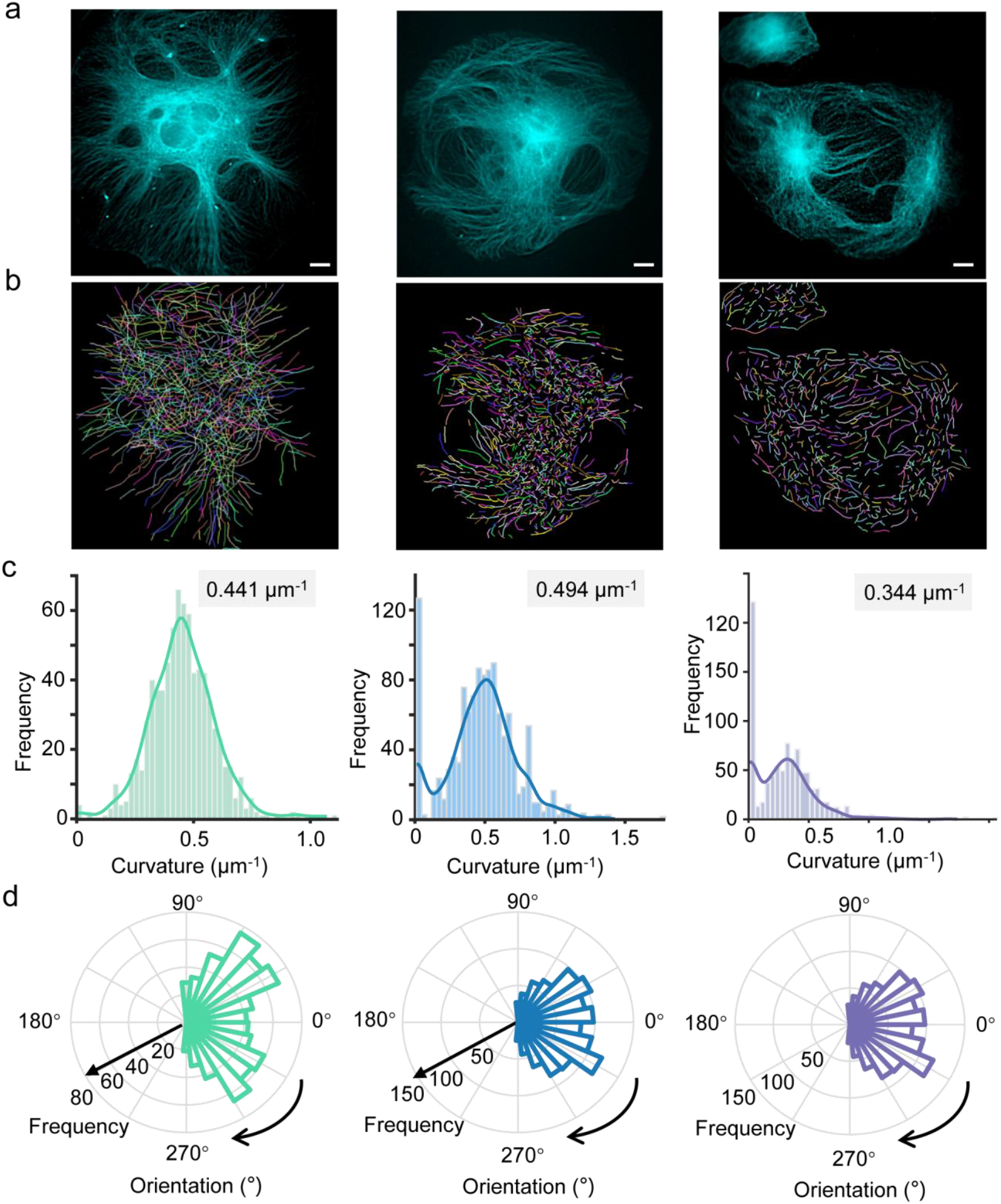
Complementary analysis with super-resolution fluorescence microscopy of intracellular microtubules. **a**, Microtubule filaments at different mitosis phases imaged under fluorescence microscopy processed by Sparse deconvolution. Scale bars: 5 µm. **b**, Segmented microtubule skeletons in ‘a’. **c**, Curvature distributions of microtubule filaments in ‘b’. **d**, Orientation analysis of microtubule filaments in ‘b’.

**Extended Data Fig. 8.**
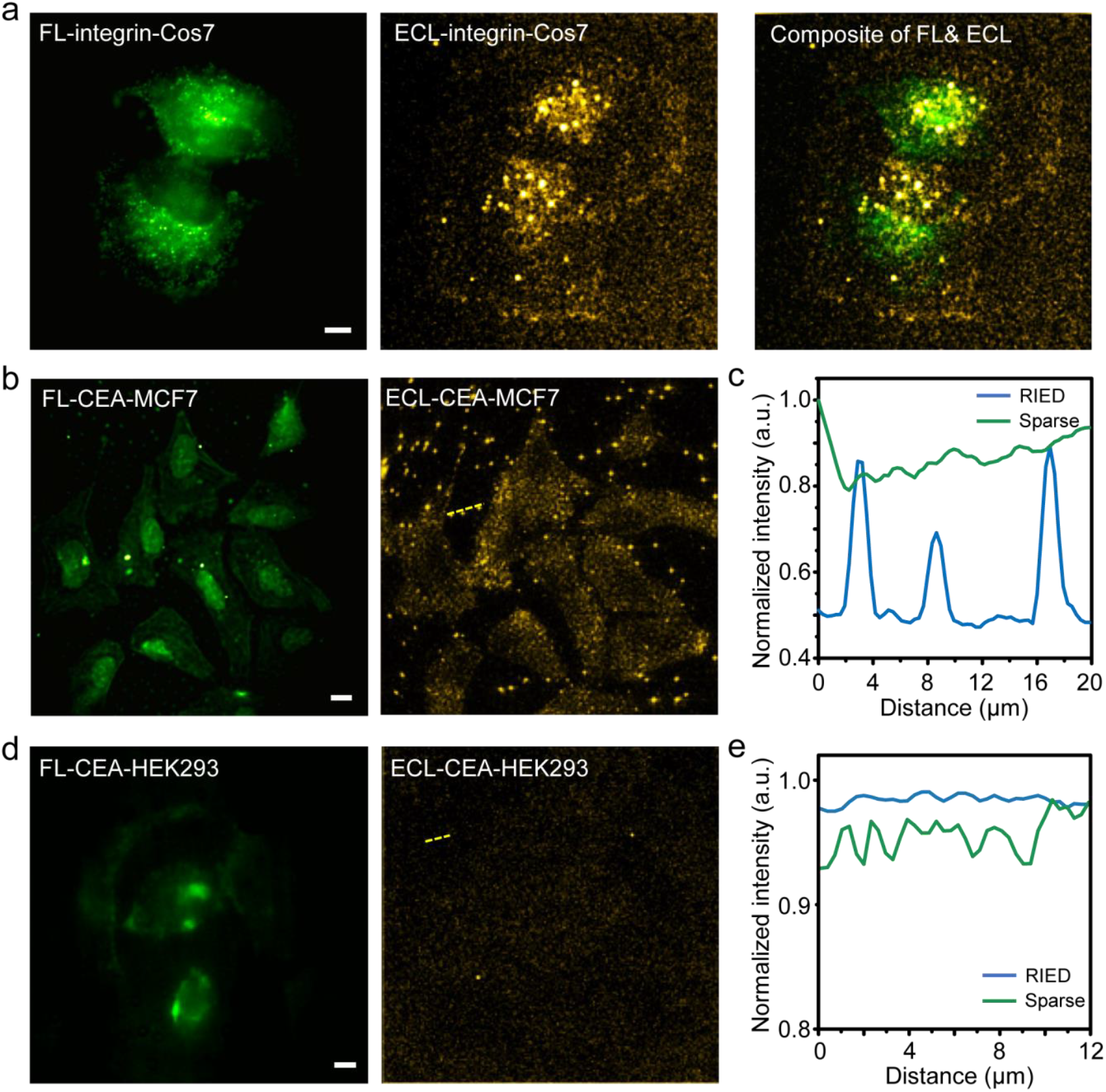
RIED for membrane protein imaging. **a**, From left to right: Sparse deconvolution reconstruction (FL, green), RIED reconstruction (ECL, yellow), and the merged results of integrin imaging in COS-7 cells. Scale bar: 5 µm. **b**, Sparse deconvolution (FL, green) and RIED (ECL, yellow) results of CEA imaging in cancer (MCF-7) cells. Scale bar: 10 µm. **c**, Intensity profiles of CEA indicated by a yellow dashed line in ‘b’. **d**, Sparse deconvolution (FL, green) and RIED (ECL, yellow) results of CEA imaging in normal (HEK293) cells. Scale bar: 10 µm. **e**, Intensity profiles of CEA indicated by a yellow dashed line in ‘d’.

